# Gut microbiome composition and predicted functions relate to growth and behavior in a Japanese preschool cohort

**DOI:** 10.64898/2026.04.15.718572

**Authors:** Shunsuke Ichikawa, Ayaka Shimura, Aoi Kikuchi, Rise Sanda, Kensaku Sasayama, Keiko Nonoue, Hiroko Tamura, Takahiro Kano, Yasuhito Shimada

## Abstract

Early childhood is a period of rapid brain maturation and gut microbiome assembly, when emerging behavioral difficulties can shape later mental health and learning trajectories. Microbiota–gut–brain communication has been implicated in neurodevelopment through microbial metabolites and immune signaling. However, most pediatric evidence comes from high-risk or clinically referred cohorts, and gut microbiome–related correlates of typical behavioral variation in community-based preschool children remain poorly defined. In a cross-sectional sample of typically developing Japanese preschool children, here we show that behavioral variation within normative ranges is associated with distinct microbiome configurations: internalizing domains cluster with signatures consistent with higher inflammatory potential and elevated nucleotide biosynthesis, whereas somatic complaints and withdrawn behavior associate with reduced respiratory and fermentative activity. Sleep-related difficulties show the broadest predicted functional footprint, including enrichment of pathways related to methyl-donor and heme biosynthesis, while externalizing domains associate with pathways involved in cell-envelope and carbohydrate remodeling. In contrast, age, height, and weight track canonical maturation of gut microbiome composition, indicating that behavioral associations are not simple proxies of growth. Together, these findings extend early-life microbiome research by resolving domain-specific associations in a low-risk Asian community sample and highlighting pathway-level candidates that may interface with neurodevelopment.

## Introduction

Preschool mental health problems are common and impairing rather than transient. Large epidemiological cohorts and a meta-analysis suggest that about 15–20% of children aged 1–7 years meet criteria for at least one mental disorder [39]. Symptoms cluster into internalizing, externalizing, and sleep or regulation problems, with sufficient consistency that preschool adaptations of standard diagnostic nosologies have been proposed [10]. Longitudinal studies show that distinct internalizing, externalizing, and co-occurring profiles are detectable by 18 months to 3 years; persistent co-occurring problems predict the greatest difficulties at school entry; and early symptom clusters, such as sleep problems at 18 months, predict increased internalizing and externalizing problems at 5 years [1, 32]. Standardized parent-report instruments are therefore central in research and practice. The Child Behavior Checklist for Ages 1½–5 (CBCL 1½–5) is widely used, provides empirically derived syndrome scales spanning internalizing, externalizing, and sleep-related problems, and shows robust factor structure and construct validity in population-based and clinical samples, including preschool children with autism spectrum disorder [1, 26]. Its use in longitudinal cohorts that track trajectories from toddlerhood into later childhood enables internationally comparable phenotyping in the present study [1, 26].

Among candidate biological markers, the gut microbiota and the microbiota–gut–brain axis have gained prominence. Intestinal microbes shape host metabolism, immune activation, and neuroendocrine signaling, and communicate with the brain via short-chain fatty acids, bile acids, tryptophan metabolites, cytokines, and vagal and enteric pathways [22]. In animal models, germ-free rearing, early antibiotic exposure, and colonization with defined microbial communities alter microglial maturation, myelination, hippocampal and prefrontal structure, stress responsivity, and anxiety- and depression-like behaviors, and several effects can be partly normalized by restoring a complex microbiota or administering psychobiotic strains [19, 33]. Fecal microbiota transplantation provides further causal evidence: microbiota from individuals with major depression, autism spectrum disorder, or attention-deficit/hyperactivity disorder can induce changes in brain structure, functional connectivity, and emotional or hyperactivity-like phenotypes in recipient mice [31, 33]. In adults, population-based metagenomics links neuroactive taxa and pathway-level features, including butyrate-producing genera such as *Faecalibacterium* and *Coprococcus* and microbial modules for γ-aminobutyric acid and dopamine metabolism, with depressive symptoms and health-related quality of life, although effects are modest and heterogeneous across cohorts [2, 28, 35]. Together, these findings motivate testing microbiota–behavior links during early childhood, when both the microbiome and neurocognitive systems mature rapidly.

Human cohort studies now extend this evidence into development from pregnancy to middle childhood, but results vary by age, population, and outcome measures. In the Barwon Infant Study, higher maternal third-trimester gut microbiota α-diversity and greater abundances of butyrate-producing Lachnospiraceae and Ruminococcaceae predicted fewer internalizing problems at 2 years of age on the CBCL [7]. In the same setting and other birth cohorts, infant gut microbiota composition and diversity during late infancy forecast CBCL internalizing problems and broader behavioral outcomes at 2–3 years, with taxa such as *Prevotella* and *Bacteroides* often implicated [17]. Longer-term community studies report that microbiota trajectories across the first 3 years relate to problem behavior and executive functions, and that patterns of diversity and composition across the first 14 years associate with internalizing, externalizing, and social anxiety symptoms around puberty [24, 41]. In typically developing preschoolers, cross-sectional studies link α-diversity, specific genera, and stool metabolomic profiles to internalizing and related dimensions; cluster-based approaches also identify configurations enriched in *Bifidobacterium* or *Bacteroides* that differ in adaptive skills and social functioning [36, 37]. Parallel work in high-risk groups suggests that very-low-birth-weight and preterm infants with early dysbiosis show altered cognitive and CBCL-based behavioral profiles, and that children with autism spectrum disorder or attention-deficit/hyperactivity disorder exhibit distinct gut microbial signatures, some of which transfer autism- or hyperactivity-like behaviors to germ-free mice [25, 31, 33]. Nevertheless, pediatric studies have largely focused on global internalizing or externalizing scores, categorical diagnoses, or selected high-risk populations, and only a few have systematically related microbial diversity, genus-level composition, and predicted functions to multiple CBCL 1½–5 syndrome scales in non-referred preschool children.

Functional resolution is also limited: although recent preschool work integrated 16S profiling with stool metabolomics to define microbiota clusters that differ in behavioral outcomes [36], predicted metabolic pathways have rarely been tested against specific domains such as anxious/depressed, withdrawn, somatic complaints, sleep problems, attention problems, and aggression. This gap matters because adult studies suggest that pathway-level features show more reproducible associations with depressive symptomatology than individual taxa [2, 28].

Most existing cohorts are from Europe or North America, and Japanese preschool children remain relatively understudied despite cultural and dietary factors that may shape both the gut microbiota and early behavior. In Japanese children aged 3–4 years, temperament assessed with the Children’s Behavior Questionnaire was associated with gut microbiota composition and diversity, and higher negative affectivity was related to lower Faecalibacterium and higher *Eggerthella* and *Flavonifractor* [34]. Another Japanese preschool study reported that children at risk for explicit emotion regulation difficulties showed higher relative abundance of *Actinomyces* and *Sutterella* than non-risk children [11]. Building on this evidence, the present study examined whether gut microbial diversity, genus-level composition, and predicted gut microbiome functions are associated with anthropometric measures and the full set of CBCL 1½–5 syndrome scales in a community sample of Japanese preschool children.

## Methods

### Ethics approval and consent to participate

The study protocol was approved by the Ethics Committee of the Faculty of Education, Mie University (approval No. 2023-02). All methods were performed in accordance with the relevant guidelines and regulations and in accordance with the Declaration of Helsinki. Written informed consent was obtained from the parents or legal guardians of all participating children.

### Participants and stool collection

To examine associations between gut microbiota, physical growth, and behavioral development in early childhood, we collected stool samples from 36 preschool children living in Mie, Okayama and Shiga Prefectures. Guardians received a stool collection kit for gut microbiota analysis (TechnoSuruga Laboratory, Shizuoka, Japan) and were instructed to collect a stool sample at home and mail it to a chemistry laboratory at the Faculty of Education, Mie University. Upon arrival at the laboratory, stool samples were immediately frozen and stored at –20 °C until DNA extraction.

### Behavioral assessment of preschool children

Guardians reported each child’s sex, date of birth, and completed the Child Behavior Checklist for Ages 1½–5 (CBCL 1½–5), a parent-report questionnaire within the Achenbach System of Empirically Based Assessment (ASEBA). The CBCL 1½–5 comprises 100 items that are rated 0 (“Not true”), 1 (“Somewhat or sometimes true”), or 2 (“Very true or often true”) and are grouped into seven empirically derived syndrome scales: Emotionally Reactive, Anxious/Depressed, Withdrawn, Somatic Complaints, Attention Problems, Aggressive Behavior, and Sleep Problems. In this study, we used the Japanese version of the CBCL 1½–5, which has demonstrated adequate reliability and validity [12]. A detailed list of questionnaire items is provided in Table S1. Item scores were summed to yield raw scores for each syndrome and the broadband Internalizing, Externalizing, and Total Problems scales. Raw syndrome scores were used both to describe normative, borderline, and clinical ranges and as continuous variables in correlation analyses, in line with Japanese CBCL 1½–5 manuals.

### Microbiome profiling and statistical analysis

Stool DNA was extracted using the NucleoSpin^®^ DNA Stool kit (Macherey-Nagel, Düren, Germany) with bead-beating. The V3–V4 region of the bacterial 16S rRNA gene was amplified and sequenced on an Illumina MiSeq platform using paired-end 2 × 300 bp reads. Reads were processed in QIIME 2 (v2024.2) with DADA2 to generate amplicon sequence variants (ASVs), and taxonomy was assigned against the EzBioCloud 16S database.

Functional profiles were predicted from ASVs using PICRUSt2 (v2.5.2) to obtain MetaCyc pathway abundances. Associations between anthropometrics, CBCL syndrome scores, and microbiome features (α-diversity indices, genus-level relative abundance, and predicted MetaCyc pathways) were evaluated using Spearman’s rank correlation. Detailed laboratory procedures and bioinformatic parameters are provided in Supplementary Methods.

## Results

### Physical characteristics and behavioral domains measured by CBCL 1½–5

A total of 36 children completed the CBCL 1½–5 (Fig. 1, Table S2). Twenty were boys and sixteen were girls. The mean age was 48.0 ± 8.2 months. Mean body size was 99.0 ± 5.2 cm in height (range 86.5–112.0 cm) and 15.6 ± 1.8 kg in weight (range 12.5–21.0 kg). Most children scored within the normative range across CBCL domains. Borderline elevations were most common for Internalizing Problems (7/36, 19.4%) and Sleep Problems (5/36, 13.9%). Clinically significant scores were observed most frequently for Externalizing Problems (6/36, 16.7%), followed by Total Problems (4/36, 11.1%). No participant met the clinical threshold for Anxious/Depressed, Withdrawn, or Attention Problems (Table S2).

**Fig. 1.**
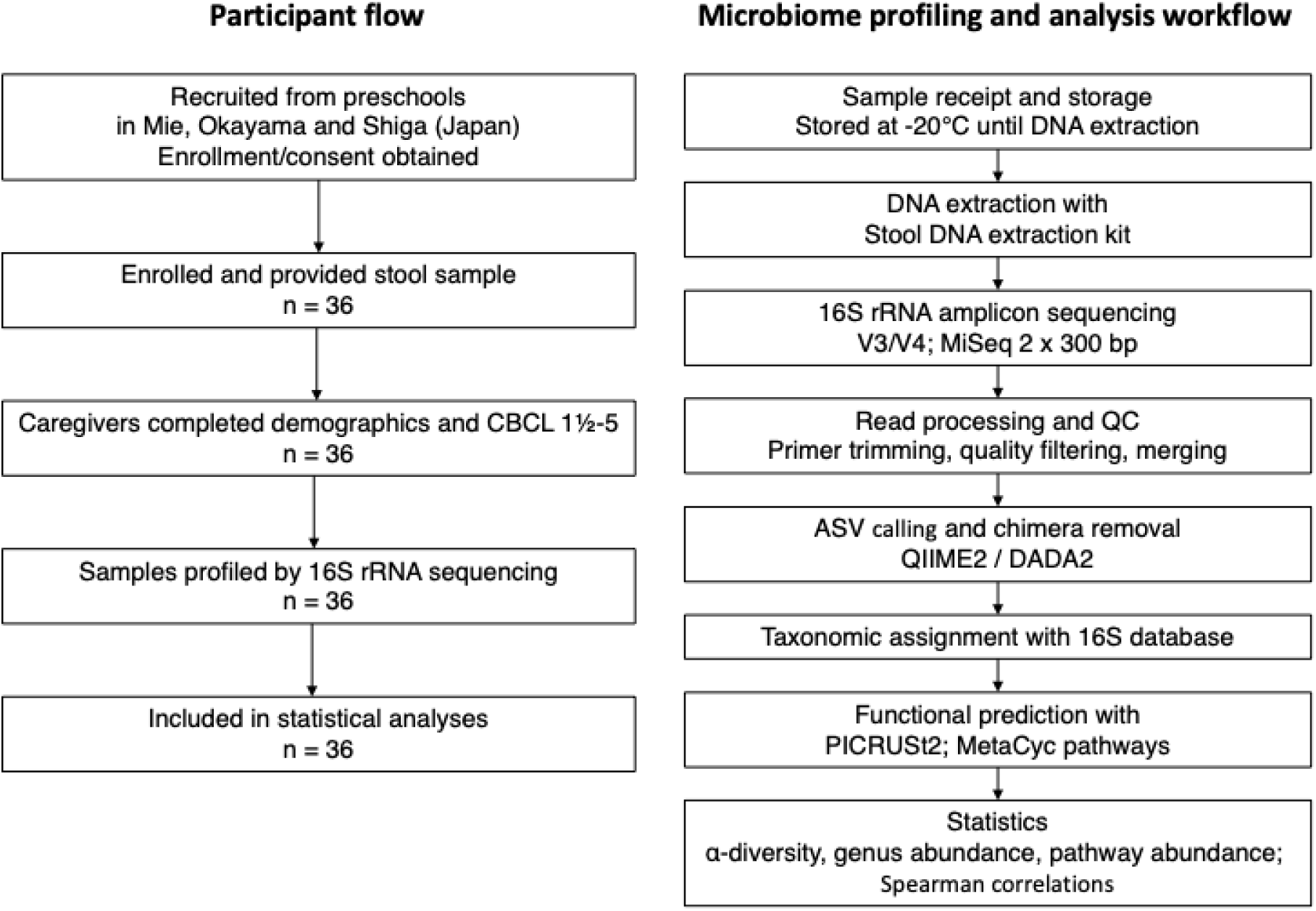
Participant flow diagram and microbiome analysis workflow

We used Spearman’s rank correlations to examine associations among age, height, weight, and CBCL domains (Fig. 2). Height and weight were strongly correlated (ρ = 0.86, p < 0.001). Age correlated positively with height (ρ = 0.60, p < 0.001) and with weight (ρ = 0.48, p = 0.003). Aggressive Behavior correlated with Emotionally Reactive (ρ = 0.59, p < 0.001) and with Anxious/Depressed (ρ = 0.55, p = 0.001), suggesting co-occurrence of aggression, emotional dysregulation, and internalizing symptoms. Emotionally Reactive correlated with Anxious/Depressed (ρ = 0.62, p < 0.001). Anxious/Depressed also correlated with Attention Problems (ρ = 0.46, p = 0.005) and Sleep Problems (ρ = 0.38, p = 0.023), suggesting shared underlying mechanisms among internalizing, attentional, and sleep domains. Withdrawn correlated with Attention Problems (ρ = 0.44, p = 0.007) and Aggressive Behavior (ρ = 0.34, p = 0.040), indicating that social withdrawal could co-occur with both attentional and externalizing features. In contrast, the Somatic Complaints scale did not demonstrate any statistically significant correlations with physical measures or other behavioral domains. Overall, CBCL domains were interrelated, whereas associations between growth indices and behavioral traits were comparatively modest.

**Fig. 2.**
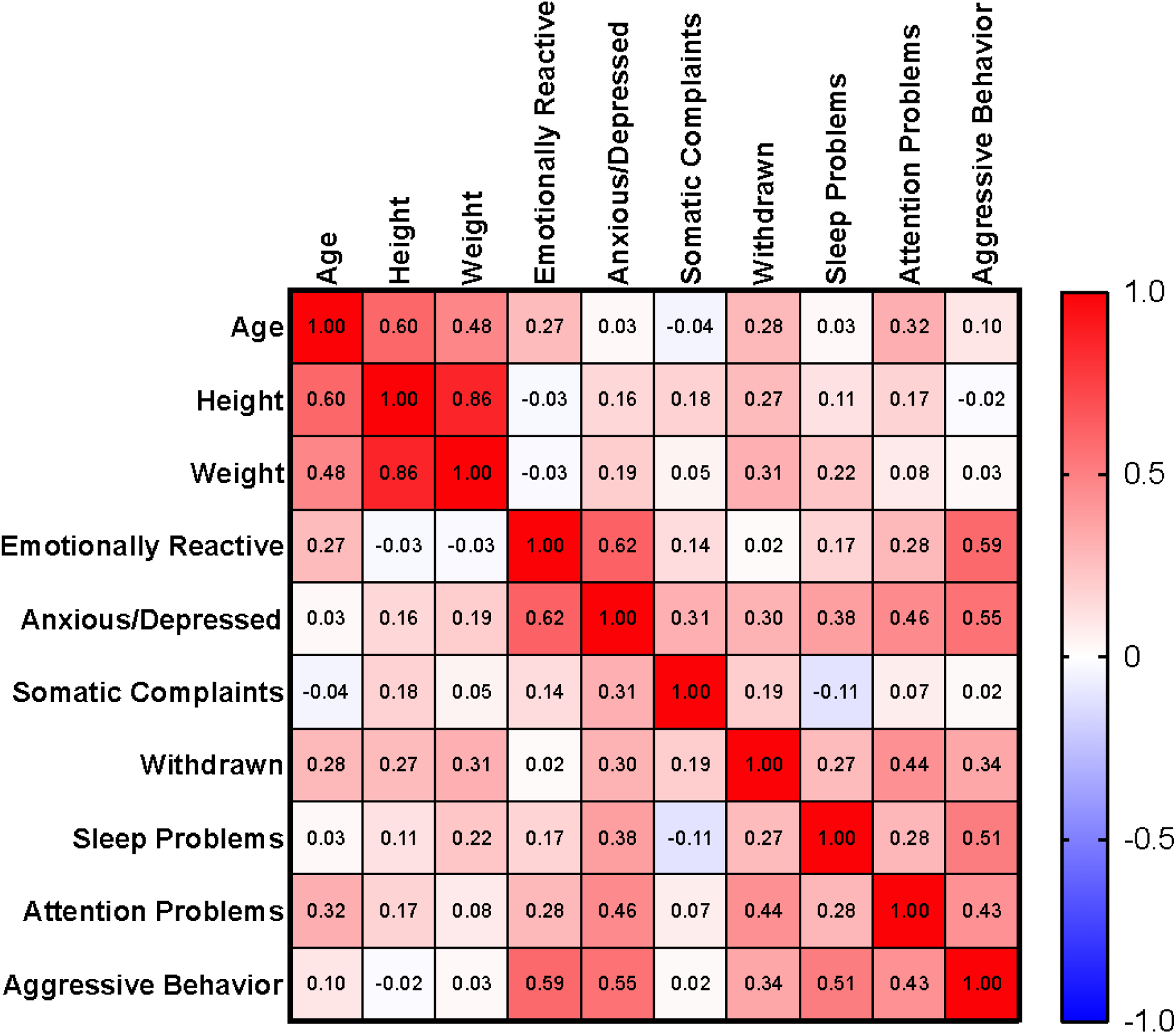
Correlation matrix of physical characteristics and CBCL 1½–5 syndrome scores in preschool children

### Specific association between attention problems and gut microbiota α-diversity

α-Diversity indices were not associated with age, height, or weight. At the nominal p < 0.05 level, Attention Problems correlated positively with observed features (ρ = 0.34, p = 0.046); Chao1 richness (ρ = 0.34, p = 0.043); and Faith’s phylogenetic diversity (ρ = 0.39, p = 0.018) (Table S3). No other syndrome scale correlated with α-diversity indices, including Emotionally Reactive, Anxious/Depressed, Somatic Complaints, Withdrawn, Sleep Problems, and Aggressive Behavior. These effects were modest and should be interpreted cautiously given the sample size, but they suggest that within-sample richness and phylogenetic diversity may align more closely with attention-related variation than with other behavioral domains or growth indicators in this cohort.

### Associations between gut microbiota and behavioral and physical development in preschool children

In total, 362 genera were identified across 36 amplicon-based microbiome profiles (Table S4). Spearman’s rank correlations between genus relative abundance and 10 traits yielded 128 nominal associations with p < 0.05 (Table S5). We primarily interpret taxa that were common, mean relative abundance ≥ 0.10%, and strongly associated with at least one trait, |ρ| ≥ 0.40, while noting additional patterns in Table 1.

**Table 1.**
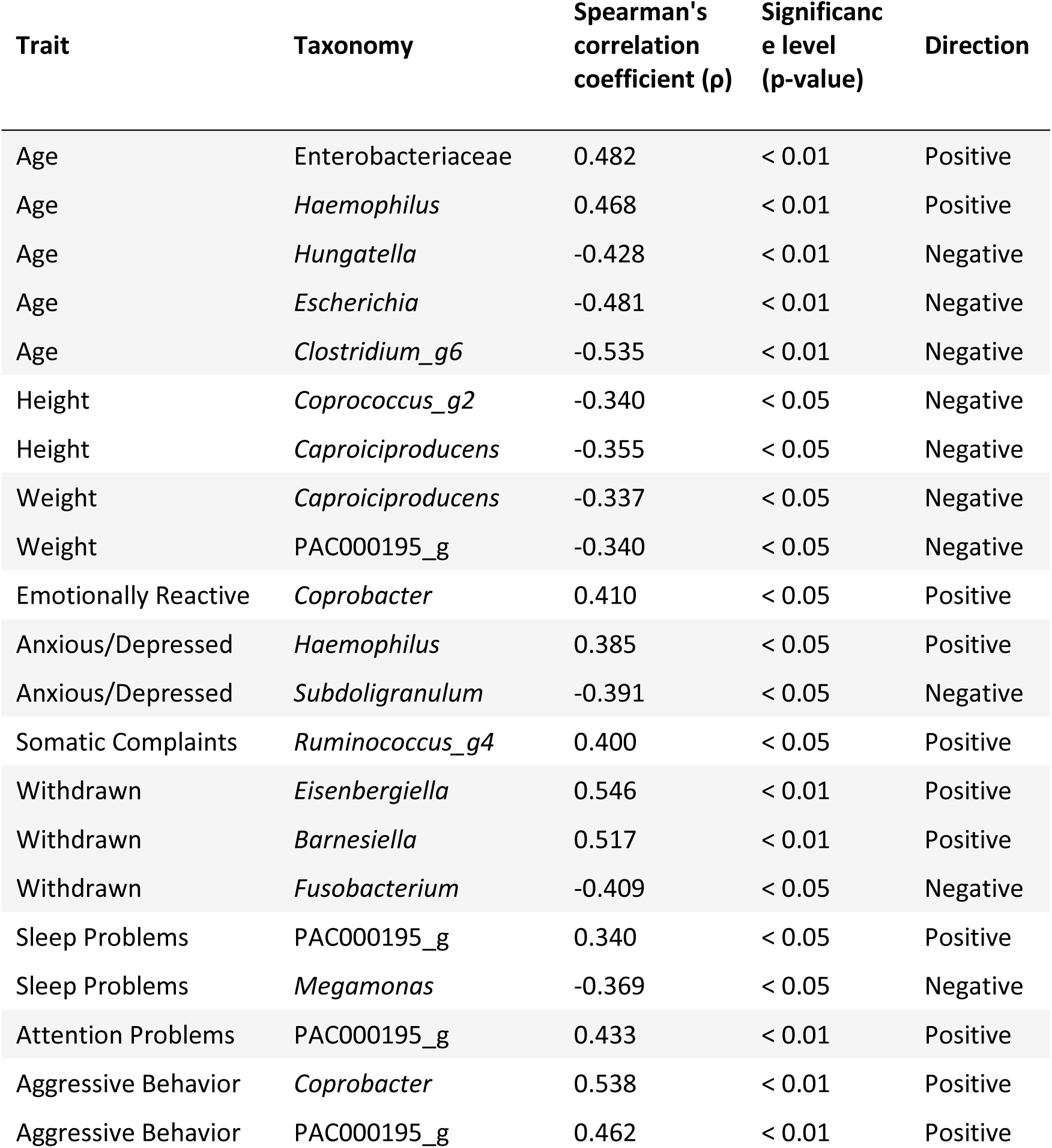
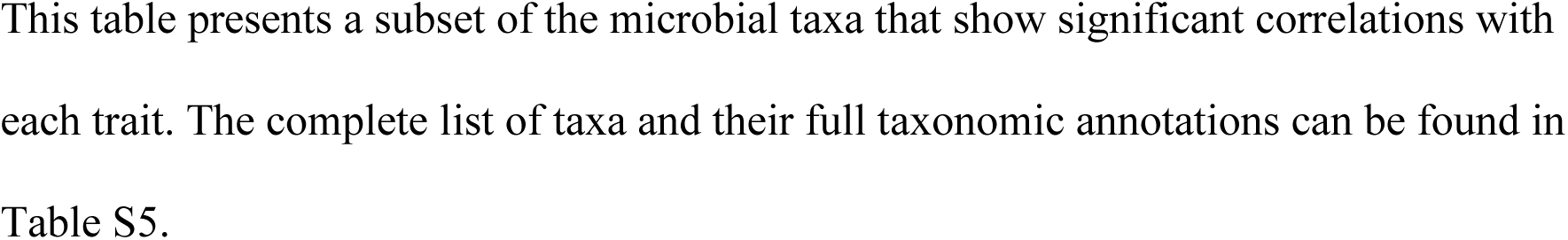
Representative gut microbial genera linked to physical characteristics and CBCL 1½–5 behavioral measures in preschool children.

| Trait | Taxonomy | Spearman's correlation coefficient ($\rho$) | Significance level (p-value) | Direction |
| --- | --- | --- | --- | --- |
| Age | Enterobacteriaceae | 0.482 | < 0.01 | Positive |
| Age | *Haemophilus* | 0.468 | < 0.01 | Positive |
| Age | *Hungatella* | -0.428 | < 0.01 | Negative |
| Age | *Escherichia* | -0.481 | < 0.01 | Negative |
| Age | *Clostridium_g6* | -0.535 | < 0.01 | Negative |
| Height | *Coprococcus_g2* | -0.340 | < 0.05 | Negative |
| Height | *Caproiciproducens* | -0.355 | < 0.05 | Negative |
| Weight | *Caproiciproducens* | -0.337 | < 0.05 | Negative |
| Weight | PAC000195_g | -0.340 | < 0.05 | Negative |
| Emotionally Reactive | *Coprobacter* | 0.410 | < 0.05 | Positive |
| Anxious/Depressed | *Haemophilus* | 0.385 | < 0.05 | Positive |
| Anxious/Depressed | *Subdoligranulum* | -0.391 | < 0.05 | Negative |
| Somatic Complaints | *Ruminococcus_g4* | 0.400 | < 0.05 | Positive |
| Withdrawn | *Eisenbergiella* | 0.546 | < 0.01 | Positive |
| Withdrawn | *Barnesiella* | 0.517 | < 0.01 | Positive |
| Withdrawn | *Fusobacterium* | -0.409 | < 0.05 | Negative |
| Sleep Problems | PAC000195_g | 0.340 | < 0.05 | Positive |
| Sleep Problems | *Megamonas* | -0.369 | < 0.05 | Negative |
| Attention Problems | PAC000195_g | 0.433 | < 0.01 | Positive |
| Aggressive Behavior | *Coprobacter* | 0.538 | < 0.01 | Positive |
| Aggressive Behavior | PAC000195_g | 0.462 | < 0.01 | Positive |

### Age

With age, *Escherichia* decreased (mean = 2.1%, ρ = –0.48, p < 0.01), and *Clostridium*_g6 also decreased (0.4%, ρ = –0.54, p < 0.01). In contrast, *Haemophilus* increased (1.3%, ρ = 0.47, p < 0.01), Enterobacteriaceae increased (0.14%, ρ = 0.48, p < 0.01) and *Hungatella* decreased (0.32%, ρ = –0.43, p < 0.01) (Table 1).

### Height and Weight

Height correlated negatively with *Coprococcus*_g2 (1.71%, ρ = –0.34, p < 0.05) and *Caproiciproducens* (0.40%, ρ = –0.36, p < 0.05). Both taxa have been described as producers of butyrate and hexanoate from complex polysaccharides [18]. Weight correlated negatively with *Caproiciproducens* (0.40%, ρ = –0.34, p < 0.05) and with PAC000195_g (0.14%, ρ = –0.34, p < 0.05). These correlations do not establish causality in this cross-sectional cohort, but they suggest lower relative abundance of these fiber-fermenting taxa in taller and heavier children.

### Anxious/Depressed

Anxious/Depressed scores were positively associated with *Haemophilus* (1.26%, ρ = 0.39, p < 0.05) and negatively with *Subdoligranulum* (0.96%, ρ = –0.39, p < 0.05). *Haemophilus* can utilize host-derived sialic acids and produce lipooligosaccharides that activate Toll-like receptor signaling [14, 15], whereas *Subdoligranulum* has been implicated in supporting mucosal barrier integrity [28, 38]. This reciprocal pattern is compatible with greater inflammatory potential and reduced barrier support in children with higher Anxious/Depressed scores, although these functional inferences remain speculative without direct measurement of host inflammatory markers or microbial metabolites.

### Somatic Complaints

Somatic Complaints were positively associated with *Ruminococcus*_g4 (1.91%, ρ = 0.40, p < 0.05), a taxon that ferments resistant starch to acetate and formate [8]. Formate has been proposed to influence sensory pathways involved in visceral pain [21], but the mechanism underlying this association remains unclear.

### Withdrawn

Withdrawn showed the strongest single effect size among the trait–taxon correlations. *Eisenbergiella* (0.63%, ρ = 0.55, p < 0.01) and *Barnesiella* (0.14%, ρ = 0.52, p < 0.01) were enriched in children with higher scores. *Eisenbergiella* can generate amino-acid-derived butyrate, and *Barnesiella* harbors bile-salt hydrolases and has been linked to modulation of host immunity and social behavior [6], suggesting potential relevance of amino-acid- and bile-acid-related microbial pathways for social withdrawal in early childhood. *Fusobacterium*, a succinate and putrescine producer, was depleted (0.23%, ρ = – 0.41, p < 0.05), supporting the view that withdrawal-related variation may involve opposing shifts across microbial metabolic routes rather than uniform changes in overall short-chain fatty acid output.

### Sleep Problems

Sleep Problems were inversely correlated with *Megamonas*, a propionate producer (1.69%, ρ = –0.37, p < 0.05), and positively correlated with PAC000195_g (0.14%, ρ = 0.34, p < 0.05). These contrasting associations raise the possibility that differences in the balance between propionate and mixed short-chain fatty acid production relate to variation in sleep regulation or circadian rhythms, although targeted metabolomic data would be needed to test this hypothesis.

### Emotional Reactivity, Attention Problems and Aggressive Behavior

Aggressive Behavior showed the highest correlations with *Coprobacter* (0.16%, ρ = 0.54, p < 0.01), and the Lachnospiraceae taxon PAC000195_g (0.14%, ρ = 0.46, p < 0.01). *Coprobacter* was also positively associated with Emotional Reactivity (0.16%, ρ = 0.41, p < 0.05) but not with Attention Problems, whereas PAC000195_g was positively associated with Attention Problems (0.14%, ρ = 0.43, p < 0.01) but not with Emotional Reactivity. *Coprobacter* belongs to a Barnesiellaceae lineage that often encodes bile salt hydrolase homologs, suggesting capacity to deconjugate primary bile acids. Deconjugated bile acids can cross the blood–brain barrier and engage central bile acid receptors in limbic circuits, and bile acid signaling has been linked to anxiety- and depression-like behaviors in rodent models [4]. These features provide one plausible route through which *Coprobacter*-enriched communities might be linked to heightened emotional lability and aggression, although such causal pathways were not tested here. PAC000195_g, in contrast, carries an extensive repertoire of carbohydrate esterases, suggesting a capacity to liberate small neuroactive metabolites from complex plant polysaccharides, but this mechanism and its potential relevance to inattention and impulsivity remain to be directly demonstrated in humans.

### Associations between gut microbiome functions and behavioral and physical development in preschool children

PICRUSt2 inference yielded 477 non-redundant MetaCyc pathways across the 36 fecal microbiome profiles. Spearman correlations between pathway relative abundance and 10 traits identified 230 nominal associations (Table S6). We primarily interpret pathways that were common (mean relative frequency ≥ 0.0010%), and at least moderately correlated with a trait (|ρ| ≥ 0.35) (Table 2).

**Table 2.** Gut microbiome functions significantly associated with physical and behavioral traits in preschool children.

| Trait | Function | Spearman's correlation coefficient ($\rho$) | Significance level (p-value) | Direction |
| --- | --- | --- | --- | --- |
| Age | L-leucine degradation I | 0.571 | < 0.01 | Positive |
| Age | adenosylcobalamin biosynthesis I (anaerobic) | 0.564 | < 0.01 | Positive |
| Age | protocatechuate degradation II (ortho-cleavage pathway) | 0.506 | < 0.01 | Positive |
| Height | L-arginine biosynthesis III (via N-acetyl-L-citrulline) | 0.356 | < 0.05 | Positive |
| Height | succinate fermentation to butanoate | -0.486 | < 0.01 | Negative |
| Weight | L-arginine biosynthesis III (via N-acetyl-L-citrulline) | 0.425 | < 0.05 | Positive |
| Weight | succinate fermentation to butanoate | -0.470 | < 0.01 | Negative |
| Emotionally Reactive | superpathway of pyrimidine deoxyribonucleotides de novo biosynthesis | 0.442 | < 0.01 | Positive |
| Emotionally Reactive | superpathway of histidine, purine, and pyrimidine biosynthesis | 0.380 | < 0.05 | Positive |
| Emotionally Reactive | allantoin degradation IV (anaerobic) | -0.361 | < 0.05 | Negative |
| Anxious/Depressed | superpathway of pyrimidine deoxyribonucleotides de novo biosynthesis | 0.440 | < 0.01 | Positive |
| Anxious/Depressed | superpathway of histidine, purine, and pyrimidine biosynthesis | 0.378 | < 0.05 | Positive |
| Somatic Complaints | aerobic respiration I (cytochrome c) | -0.401 | < 0.05 | Negative |
| Somatic Complaints | cob(II)yrinate a,c-diamide biosynthesis I (early cobalt insertion) | -0.395 | < 0.05 | Negative |
| Somatic Complaints | L-tyrosine degradation I | -0.387 | < 0.05 | Negative |
| Withdrawn | superpathway of 2,3-butanediol biosynthesis | -0.433 | < 0.01 | Negative |
| Withdrawn | heterolactic fermentation | -0.409 | < 0.05 | Negative |
| Withdrawn | Bifidobacterium shunt | -0.406 | < 0.05 | Negative |
| Sleep Problems | L-methionine biosynthesis III | 0.473 | < 0.01 | Positive |
| Sleep Problems | L-methionine biosynthesis I | 0.470 | < 0.01 | Positive |
| Sleep Problems | superpathway of S-adenosyl-L-methionine biosynthesis | 0.463 | < 0.01 | Positive |
| Sleep Problems | superpathway of heme b biosynthesis from uroporphyrinogen-III | 0.463 | < 0.01 | Positive |
| Attention Problems | GDP-D-glycero-α-D-manno-heptose biosynthesis | 0.434 | < 0.01 | Positive |
| Attention Problems | superpathway of fucose and rhamnose degradation | -0.497 | < 0.01 | Negative |
| Attention Problems | L-fucose degradation I | -0.435 | < 0.01 | Negative |
| Aggressive Behavior | superpathway of heme b biosynthesis from uroporphyrinogen-III | 0.461 | < 0.01 | Positive |
| Aggressive Behavior | tRNA processing | 0.404 | < 0.05 | Positive |
| Aggressive Behavior | ADP-L-glycero-β-D-manno-heptose biosynthesis | 0.379 | < 0.05 | Positive |

### Age

Older preschool children showed increased predicted capacities for branched-chain amino-acid catabolism and cofactor production. The strongest signal was L-leucine degradation I (LEU-DEG2-PWY) (0.0043% mean relative frequency, ρ = 0.57, p < 0.01), followed by adenosylcobalamin biosynthesis I (PWY-5507) (0.0190%, ρ = 0.56, p < 0.01). Leucine degradation yields acetyl-CoA and acetoacetate, whereas adenosylcobalamin biosynthesis suggests increased predicted capacity for vitamin B_12_ production. Together, these patterns are consistent with enhanced microbial energy harvest and cobalamin-dependent metabolism with age. Protocatechuate degradation II (PROTOCATECHUATE-ORTHO-CLEAVAGE-PWY) (0.0019%, ρ = 0.51, p < 0.01) also increased with age, indicating a parallel expansion of polyphenol breakdown as diets diversify.

### Height and Weight

Height was positively linked to amino-acid anabolism rather than fermentation. L-arginine biosynthesis III via N-acetyl-L-citrulline (PWY-5154), correlated positively (0.360%, ρ = 0.36, p < 0.05), whereas succinate fermentation to butanoate (PWY-5677) was inversely related (0.0322%, ρ = –0.49, p < 0.01). Taller children therefore tended to host microbiota with higher predicted capacity for arginine biosynthesis and lower inferred reliance on reductive succinate pathways. Body weight reproduced the height pattern for arginine biosynthesis (0.360%, ρ = 0.43, p < 0.05), and for succinate fermentation to butanoate (0.0322%, ρ = –0.47, p < 0.01). These associations suggest coordinated adjustments in predicted amino-acid and energy metabolism along gradients of somatic growth in this cohort.

### Emotionally Reactive

Higher Emotional Reactivity scores coincided with enrichment of pathways related to nucleotide biosynthesis pathways. The top pathway was superpathway of pyrimidine deoxyribonucleotide *de novo* biosynthesis (PWY-7211) (0.287%, ρ = 0.44, p < 0.01), followed by superpathway of histidine, purine, and pyrimidine biosynthesis (PRPP-PWY) (0.315%, ρ = 0.38, p < 0.05). These associations were accompanied by a negative link to allantoin degradation IV (PWY0-41) (0.0078%, ρ = –0.36, p < 0.05). Together, this pattern is consistent with microbiota enriched in *de novo* nucleotide production and altered nitrogen salvage in children with higher emotional reactivity, but the functional implications for host neurotransmitter precursors remain speculative without direct metabolomic data.

### Anxious/Depressed

Anxious/Depressed scores showed a similar functional profile, with positive correlations to superpathway of pyrimidine deoxyribonucleotides *de novo* biosynthesis (PWY-7211) (0.287%, ρ = 0.44, p < 0.05) and superpathway of histidine, purine, and pyrimidine biosynthesis (PRPP-PWY) (0.315%, ρ = 0.38, p < 0.05). No negatively correlated high-frequency pathways surpassed the thresholds, suggesting that enhanced predicted *de novo* nucleotide synthesis was the dominant shift accompanying anxious traits in this sample.

### Somatic Complaints

Somatic Complaints were inversely associated with aerobic respiration I (cytochrome c) (PWY-3781) (0.0402%, ρ = –0.40, p < 0.05), cob(II)yrinate a,c-diamide biosynthesis I (early cobalt insertion) (PWY-7377) (0.145%, ρ = –0.40, p < 0.05) and L-tyrosine degradation I (TYRFUMCAT-PWY) (0.0014%, ρ = –0.39, p < 0.05). These signals indicate reduced predicted respiratory capacity and aromatic amino-acid catabolism in children reporting more somatic symptoms.

### Withdrawn

Withdrawn was most strongly associated with lower predicted fermentative capacity. Three high-frequency fermentation routes were inversely correlated, including superpathway of 2,3-butanediol biosynthesis (PWY-6396) (0.0151%, ρ = –0.43, p < 0.01), heterolactic fermentation (P122-PWY) (0.0818%, ρ = –0.41, p < 0.05), and the *Bifidobacterium* shunt (P124-PWY) (0.0976%, ρ = –0.41, p < 0.05). This pattern suggests reduced representation of classical bifidobacterial and lactic fermentative pathways in children with higher withdrawn scores.

### Sleep Problems

Sleep disturbance displayed the broadest functional footprint, dominated by sulfur-containing amino-acid and heme biosynthesis. Positive correlations spanned L-methionine biosynthesis III (HSERMETANA-PWY) (0.313%, ρ = 0.47, p < 0.01), L-methionine biosynthesis I (HOMOSER-METSYN-PWY) (0.283%, ρ = 0.47, p < 0.01), superpathway of S-adenosyl-L-methionine biosynthesis (MET-SAM-PWY) (0.372%, ρ = 0.46, p < 0.01), and superpathway of heme *b* biosynthesis from uroporphyrinogen-III (PWY0-1415) (0.0442%, ρ = 0.46, p < 0.01), suggesting a potential link between microbial methyl-group turnover, heme production, and sleep quality in preschool children.

### Attention Problems

Higher Attention Problems scores were marked by a striking depletion of fucose and rhamnose catabolism: superpathway of fucose and rhamnose degradation (FUC-RHAMCAT-PWY) (0.211%, ρ = –0.50, p < 0.01) and L-fucose degradation I (FUCCAT-PWY) (0.157%, ρ = –0.44, p < 0.01) were both negatively associated. Conversely, GDP-D-glycero-α-D-manno-heptose biosynthesis (PWY-6478) (0.0931%, ρ = 0.43, p < 0.01) was enriched. Together, these shifts may reflect altered microbial handling of host-derived glycans and increased heptose-nucleotide sugar biosynthesis, suggesting changes in bacterial cell-surface glycoconjugate metabolism in children with higher attention problem scores.

### Aggressive Behavior

Aggressive Behavior correlated positively with porphyrin metabolism and core RNA handling. Key pathways included superpathway of heme *b* biosynthesis from uroporphyrinogen-III (PWY0-1415) (0.0442%, ρ = 0.46, p < 0.01), tRNA processing (PWY0-1479) (0.0981%, ρ = 0.40, p < 0.05), and ADP-L-glycero-β-D-manno-heptose biosynthesis (PWY0-1241) (0.0736%, ρ = 0.38, p < 0.05). These functions support intensified redox activity and cell-wall maturation, a pattern compatible with microbiota that have relatively high predicted biosynthetic and growth potential in children with higher aggression scores.

## Discussion

### Gut microbiota along physical growth trajectories

Age, height, and weight were strongly intercorrelated, but their correlations with CBCL domain scores were modest (Fig. 2), suggesting that behavioral variation was not a simple proxy for somatic development. Somatic indices nevertheless tracked canonical microbiota maturation and predicted functions. The decline of Escherichia and other facultative anaerobes with age, together with increasing representation of strictly anaerobic genera, mirrors cohort data describing the transition toward a more stable, adult-like microbiota during the preschool years [23]. Age-related enrichment of leucine degradation and adenosylcobalamin biosynthesis suggests increasing microbial contributions to branched-chain amino-acid catabolism and vitamin B_12_ availability as diets diversify. Height and weight were positively associated with arginine biosynthesis and inversely associated with succinate-to-butanoate fermentation, pointing to a shift away from reductive succinate pathways in larger children. The weak coupling between growth indices and CBCL traits supports the interpretation that the domain-specific signatures below are unlikely to be simple proxies of maturation.

### Overview of microbiota–behavior associations in a community sample

In this cross-sectional study of typically developing Japanese preschool children, gut microbial community structure and predicted functions showed coordinated associations with multiple CBCL 1½–5 domains. Even within normative ranges, variation in internalizing, externalizing, and sleep-related difficulties aligned with distinct taxonomic and metabolic patterns. This extends early-life microbiota–gut–brain research beyond high-risk or clinically referred cohorts and is broadly consistent with longitudinal links between infant microbiota and later internalizing and externalizing problems, executive function, and temperament [17, 23, 41]. In our data, anxious/depressed symptoms and emotional reactivity clustered with taxa and pathways suggestive of higher inflammatory potential and greater biosynthetic activity, whereas somatic complaints and withdrawn behavior aligned with reduced respiratory and fermentative activity. This pattern resonates with preschool studies implicating microbial metabolic output, including short-chain fatty acid profiles, rather than diversity alone [36, 37].

### Attention and aggression: distinct externalizing–microbiota relationships

Attention problem scores were positively associated with α-diversity indices, including observed features, Chao1 richness, and Faith’s phylogenetic diversity. Case–control studies in children and adolescents with clinically diagnosed attention-deficit/hyperactivity disorder (ADHD) variably report reduced α-diversity or no consistent differences [3, 29].

Our community sample captures dimensional variation rather than categorical diagnoses and suggests that higher richness can co-occur with more pronounced attentional difficulties in the preschool period. A similar direction has been reported in another Japanese preschool cohort, where higher gut microbiota diversity was associated with greater impulsivity [34]. These patterns motivate longitudinal cohorts that integrate dietary and medication histories to clarify whether α-diversity reflects transient development, confounding, or early risk.

Taxonomic and functional profiles further linked attention problems and aggressive behavior to genera with plausible neuroactive potential. *Coprobacter* and the Lachnospiraceae taxon PAC000195_g, which correlated with aggressive behavior and attention problems respectively, encode bile-salt hydrolases and carbohydrate esterases, suggesting an enhanced capacity to remodel host bile acids and complex polysaccharides. Such remodeling may increase unconjugated bile acids and other small products that influence limbic excitability and prefrontal control circuits, consistent with animal-model evidence linking bile acid and microbial metabolite signaling to hyperactivity and impulsivity [9, 33]. Aggressive Behavior was associated with ADP-L-glycero-β-D-manno-heptose biosynthesis (PWY0-1241). This pathway generates ADP-heptose, an inner-core lipopolysaccharide precursor that can activate intracellular innate immune signaling and elevate inflammatory tone [13].

### Microbial signatures of internalizing difficulties in preschool children

Internalizing traits showed a complementary microbial profile. Anxious/depressed scores were positively associated with *Haemophilus* and negatively associated with *Subdoligranulum*, a sentinel butyrate producer linked to epithelial barrier support and inversely associated with depressive symptoms in adults [16, 28]. *Haemophilus* can exploit host sialic acids and produce lipooligosaccharides that activate Toll-like receptor 4, promoting mucosal inflammation and systemic immune activation [5, 14]. The reciprocal shift therefore supports a model in which higher anxious/depressed scores align with higher inflammatory potential and reduced barrier-supportive capacity. This interpretation is consistent with Japanese preschool findings linking emotion-regulation risk to higher *Actinomyces* and *Sutterella* [11]. Functionally, these traits aligned with enrichment of *de novo* pyrimidine and broader nucleotide biosynthesis pathways, compatible with more biosynthetically active communities that may influence host tryptophan, purine, and pyrimidine pools relevant to stress-response circuitry [9, 27].

Somatic complaints were associated with reduced inferred aerobic respiration and tyrosine degradation, suggesting lower respiratory flexibility and diminished aromatic amino-acid catabolism. Because aromatic amino-acid-derived metabolites, including tyrosine-derived amines, have been linked to visceral pain hypersensitivity and autonomic regulation, reduced microbial engagement with these substrates may contribute to altered interoceptive signaling [42]. Withdrawn behavior, in contrast, was associated with enrichment of *Eisenbergiella* and *Barnesiella* and depletion of *Fusobacterium*, together with reduced representation of classical fermentative pathways such as the *Bifidobacterium* shunt and heterolactic fermentation. These findings are consistent with reports linking lower bifidobacterial activity and altered amino-acid fermentation to social withdrawal and suggest that shifts in amino-acid-derived short-chain fatty acids may be relevant for social engagement in early childhood [30].

### Microbiome functional pathways associated with sleep problems

Sleep problems showed the broadest functional footprint, with positive correlations to microbial methionine and S-adenosyl-methionine biosynthesis and to heme production pathways. These processes regulate methyl-group availability and redox-active cofactor synthesis, which can influence circadian clock machinery, melatonin production, and oxidative stress responses. Recent studies link sleep duration and efficiency in preschoolers to microbiota profiles and to microbial genes involved in amino-acid and cofactor metabolism [20, 40]. Our data add that sleep disturbance aligns with enrichment of sulfur-containing amino-acid biosynthesis and porphyrin pathways, supporting pathway-level candidates for future mechanistic and interventional work.

### Methodological considerations and future directions

These findings should be interpreted as associative rather than causal. The modest sample size limits statistical power and may reduce generalizability beyond Japanese community-dwelling preschool children, and the cross-sectional design cannot determine whether microbiome differences precede behavioral traits, reflect shared environments, or arise through bidirectional feedback. Larger longitudinal cohorts with repeated sampling and direct multi-omic measurements will be required to test directionality and mechanism.

Across domains, signals converged on microbial capacities related to nucleotide synthesis, amino-acid and methyl-group metabolism, bile-acid and glycan transformation, and heptose-containing lipopolysaccharide biosynthesis. These pathways plausibly interface with neurodevelopment by shaping metabolite availability, epithelial and immune tone, and neuroendocrine signaling relevant to emotional regulation, attention, and sleep. Future work that combines repeated stool sampling with detailed diet and medication histories and concurrent measurement of relevant metabolites and host markers can test whether microbial methyl-donor metabolism, bile-acid remodeling, and glycan utilization mediate changes in sleep and behavior over time. If replicated, these signatures could support microbiome-informed prevention studies in early childhood, including pragmatic trials of dietary modulation and targeted prebiotic or probiotic approaches, with outcomes anchored to standardized behavioral scales and objective sleep measures. Over the longer term, microbiome-based markers could complement psychosocial and educational strategies by adding a biologically grounded component to early risk monitoring and support.

## Statements and Declarations

### Ethics approval

The study was approved by the ethics committee in Faculty of Education, Mie University (No. 2023-02).

### Consent to participate

Written informed consent was obtained from the children’s guardians.

### Consent to publish

Not applicable. This manuscript does not contain any individual person’s identifiable data.

### Data availability

Raw 16S rRNA gene sequencing data and associated metadata have been deposited in the NCBI Sequence Read Archive (SRA) under accession number PRJNA1392820. Processed feature tables, taxonomic assignments, and analysis scripts are available from the corresponding author upon reasonable request.

## Abbreviations

CBCL 1½–5: Child Behavior Checklist for ages 1½–5
ASEBA: Achenbach System of Empirically Based Assessment
ASV: Amplicon Sequence Variant
EC: Enzyme Commission
KEGG: Kyoto Encyclopedia of Genes and Genomes
KO: KEGG Ortholog
COG: Clusters of Orthologous Groups
PICRUSt2: Phylogenetic Investigation of Communities by Reconstruction of Unobserved States 2
QIIME 2: Quantitative Insights Into Microbial Ecology 2
SCFA: Short-Chain Fatty Acids

## Acknowledgements

We sincerely thank the children, parents, and teachers at participating kindergartens and nursery schools for their cooperation with fecal sample collection and questionnaire completion.

## Author contributions

S.I. and Y.S. conceptualized and designed the study. S.I., K.S., K.N., H.T., and T.K. collected the data. S.I., A.S., A.K., R.S., and Y.S. analyzed and interpreted the data. S.I. and Y.S. drafted and revised the manuscript. All authors reviewed and approved the final manuscript for submission.

