## Supplementary methods for "Gut microbiome composition and predicted functions relate to growth and behavior in a Japanese preschool cohort"

**DNA extraction from stool**
First, 220 µL of the fecal suspension solution was transferred to MN Bead Tubes type A (Macherey-Nagel, Düren, Germany). Then, 850 µL of Buffer ST1 in NucleoSpin^®^ DNA Stool kit (Macherey-Nagel, Düren, Germany) was added, and the tubes were shaken horizontally for two to three seconds to suspend the sample. After incubation at 70°C for five minutes, each bead tube was placed in the Homogenizer ShakeMan 6 (Bio Medical Science, Tokyo, Japan) and homogenized for a total of 10 minutes at room temperature (six cycles of 90 seconds at 2,500 rpm, with 30-second intervals). The tubes were then centrifuged at 13,000 × g for three minutes, and 600 µL of the supernatant was transferred to a new 2-mL tube. Total DNA of gut bacterial origin was extracted from the stool samples using the NucleoSpin^®^ DNA Stool kit.

**Bacterial community analysis**
Using the extracted stool DNA as a template, the V3/V4 region of the 16S rRNA gene was amplified by PCR with the primers F (ACACTCTTTCCCTACACGACGCTCTTCCGATCT-NNNNN–CCTACGGGNGGCWGCAG) and R (GTGACTGGAGTTCAGACGTGTGCTCTTCCGATCT-NNNNN–GACTACHVGGGTATCTAATCC). KOD One^®^ PCR Master Mix (Toyobo, Osaka, Japan) was used under the following conditions: denaturation at 98°C for 10 seconds, annealing at 55°C for 5 seconds, extension at 68°C for 15 seconds, for a total of 25 cycles. The amplified PCR products were purified with VAHTS DNA Clean Beads (Vazyme). A sequencing library was then prepared, and paired-end sequencing (2 × 300 bp) was performed using the MiSeq System and the MiSeq Reagent Kit v3 (Illumina).

After sequencing, reads that perfectly matched the primer sequences at the beginning of the reads were extracted using the fastx_barcode_splitter tool in the FASTX-Toolkit (ver. 0.0.14). In cases where primer sequences included N-mix (total of 36 patterns, i.e., 6 forward × 6 reverse), this procedure was repeated accordingly. The primer sequences were then trimmed using fastx_trimmer in the FASTX-Toolkit. Next, sequences with a quality score below 20 were removed using Sickle (ver. 1.33), and any reads shorter than 130 bases (and their paired reads) were discarded. Paired-end reads were merged with FLASH (ver. 1.2.11). Chimeric and noisy sequences were removed using the DADA2 plugin in Qiime2 (ver. 2024.2), which then generated representative sequences and an ASV table. Taxonomic assignment was performed by comparing these representative sequences to the EzBioCloud 16S database using the feature-classifier plugin in Qiime2. Using the frequency table and representative sequences generated by Qiime2, functional profiles were constructed with PICRUSt2 (ver. 2.5.2) based on enzyme commission numbers (EC), KEGG orthologs (KO), Clusters of Orthologous Groups (COG), and MetaCyc pathways. Spearman’s rank correlation coefficients were then calculated to examine associations between anthropometric measures and each CBCL 1½–5 syndrome scale and gut microbiota features, including α-diversity indices, genus-level relative abundances, and PICRUSt2-predicted MetaCyc pathways.
